# Modelling the probability of meeting IUCN Red List criteria to support reassessments

**DOI:** 10.1101/2023.06.08.544254

**Authors:** Etienne Georges Henry, Luca Santini, Stuart Butchart, Manuela Gonzalez-Suarez, Pablo Miguel Lucas, Ana Benitez-Lopez, Giordano Mancini, Martin Jung, Pedro Cardoso, Alexander Zizka, Carsten Meyer, H. Resit Akcakaya, Alex Berryman, Victor Cazalis, Moreno Di Marco

## Abstract

Comparative extinction risk analysis - which predicts species extinction risk from correlation with traits or geographical characteristics - has gained research attention as a promising tool to support extinction risk assessment in the IUCN Red List of Threatened Species. However, its uptake has been very limited so far, possibly because these models only predict a species’ Red List category, without indicating which Red List criteria may be triggered by which such approaches cannot easily be used in Red List assessments. We overcome this implementation gap by developing models that predict the probability of species meeting individual Red List criteria. Using data on the world’s birds, we evaluated the predictive performance of our criterion-specific models and compared it with the typical criterion-blind modelling approach. We compiled data on biological traits (e.g., range size, clutch size) and external drivers (e.g., change in canopy cover) often associated with extinction risk. For each specific criterion, we modelled the relationship between extinction risk predictors and species’ Red List category under that criterion using ordinal regression models. We found criterion-specific models were better at predicting threatened species compared to a criterion-blind model (higher sensitivity), but less good at predicting not threatened species (lower specificity). As expected, different covariates were important for predicting threat status under different criteria, for example change in annual temperature was important to predict criteria related to population trends, while clutch size was important for criteria related to restricted area of occupancy or small population size. Our criteria-specific method can support Red List assessors by producing outputs that identify species likely to meet specific criteria, and which are the most important predictors: these species can be prioritised for re-evaluation. We expect this new approach to increase the uptake of extinction risk models in Red List assessments, bridging a long-standing research-implementation gap.

## Introduction

Over recent decades, the IUCN Red List of Threatened species (henceforth “Red List”) has become the global standard for species’ extinction risk assessments (Mace et al., 2008; Rodrigues et al., 2006; Betts et al., 2020). A Red List assessment is based on five complementary criteria with quantitative thresholds relating to population and distribution size, structure and trends, to assign species to categories of extinction risk (IUCN, 2012; Mace et al., 2008). The Red List now includes assessments for over 150,000 species of animals, fungi and plants (IUCN, 2022) but, despite its great importance for conservation action and policy (Betts et al., 2020; Williams et al., 2021; Hoffmann et al., 2010), it is insufficiently funded (Juffe-Bignoli et al., 2016). As a consequence, the Red List faces important challenges in keeping assessments up to date (i.e., <10 years old), and reducing the proportion of Data Deficient species (Rondinini et al., 2014; Cazalis et al., 2022).

Different methods have been developed to support Red List assessments and address the above challenges (e.g., Buchanan et al. 2008; Santini et al. 2019; Bachman et al. 2019; Cazalis et al. 2023; see Cazalis et al. 2022 for an overview). Among them, comparative extinction risk models link extinction risk (i.e., Red List categories) with species’ biological traits (e.g., body mass, habitat specialisation, range size) and external drivers of risk (e.g., human density, land-use change, climate change; Purvis et al. 2000, Chichorro et al., 2019, 2022). The models are then used to predict the Red List categories of species that have not yet been assessed for the Red List (Zizka et al., 2021b, Darrah et al., 2017), that are Data Deficient (Luiz et al., 2016, Bland and Böhm 2016), or that need updating (Lucas et al. 2023, Di Marco et al., 2014), with the objective of accelerating and helping to prioritise the work of Red List assessors by providing additional information on species’ extinction risk.

These comparative extinction risk models predict Red List categories met under any of the five possible criteria (based on e.g., distribution, abundance, population trends) thus ignoring potential differences in their driving forces. This “criterion-blind” approach assumes all criteria can be predicted from the same covariates, while in reality the criteria address different sources of risk that are spatially structured (Figure S1) and some are likely easier to predict than others. For instance, predicting species threatened under criterion B1 (restricted extent of occurrence combined with some subcriteria) is relatively easy using range size, while it may be harder to find relevant covariates to predict species threatened under criterion A3 (future population decline). This suggests that species classified under different criteria might have different risk correlates and face different prediction uncertainties, which might have contributed to the low performance of some criterion-blind models when tested on independent samples of species (Di Marco, 2022). In some cases, a covariate could even have opposite effects for different criteria. For instance, species with large body mass tend to have low population density (Silva and Downing, 1995; Santini et al. 2018) and might be more likely to trigger criterion C1 (small population size and decline), but such species tend to have large ranges (Newsome et al., 2020) hence are less likely to trigger criterion B. Importantly, ignoring the diversity of criteria limits the uptake of comparative extinction risk models by assessors, who need criteria-specific information (Cardillo and Meijaard, 2012; Cazalis et al., 2022).

Here we aim to overcome this research-implementation gap by developing criterion-specific extinction risk models, and comparing their performance and applicability to a classic criterion-blind model. While the latter estimates the probability of a species to be threatened under any criterion, our criterion-specific model estimates such probability independently for each individual criterion (A1, A2, A3, etc). While benefiting from the power of the multi-species comparisons, this approach better encompasses the diversity of reasons that may qualify a species as threatened in the Red List, and provides assessors with an output that is directly related with the information needed for assessments. We use birds as a study group to test our approach as they are the most consistently assessed group across Red List criteria (Cazalis et al., 2022), with very few Data Deficient species (0.4%), they have been used in many previous comparative extinction risk analyses (e.g., Olah et al., 2018; Richards et al., 2021; White and Bennett, 2015), and they present great variation in their response to human pressure (Lees et al., 2022, Cazalis, 2022).

## Methods

We compiled data on avian biological traits associated with extinction risk (e.g., range size, clutch size) as well as external risk drivers operating within species ranges (e.g., change in forest canopy cover, distance to cities). We modelled the relationship between extinction risk predictors and each species’ Red List category met under each specific criterion, using ordinal regression models which best match the ordinal structure of the Red List categories (Lucas et al., 2019, 2023; Luiz et al., 2016), and combined these models into a single final prediction. We then compared the performance of these criterion-specific models with a criterion-blind approach, as well as the importance of different predictors in each approach. Finally, we evaluated the potential conservation applications of the criterion-specific approach. All analyses were conducted in R version 4.0.2 (R Core Team, 2020).

### 1. Extinction risk predictors

For each of the 11,162 bird species included in the Red List version 2021-3 (BirdLife International, 2022), we gathered information on species biological traits that are associated with extinction risk (e.g., Tobias and Pigot 2019; Olah et al. 2018; Ripple et al. 2017) considering five types (see details and rationale in Table S1): morphological (body mass, beak length, hand-wing index), behavioural (nocturnality, migratory status), life history traits (clutch size, generation length), ecological (trophic niche, forest dependency, habitat breadth) and geographical (insularity, range size). However, species’ intrinsic traits alone cannot predict extinction risk (González-Suárez et al., 2013; Chichorro et al., 2022) and it is key to include measurements of the impact of human activities within the species’ range (Di Marco et al., 2014; Murray et al., 2014). We thus also included proxies for habitat alteration and degradation (extent and change of cropland and forest cover), human encroachment (human density and trends, proportion of rural population, travel time to the nearest city) and past and contemporary climate change (difference in precipitation and temperature) within each species range (see details in Table S1).

### A. Species traits and characteristics

We used the distribution maps published in BirdLife International and Handbook of the Birds of the World (2021), filtering polygons with high probability of presence (‘extant’) and of ‘native’ origin during the breeding season (‘resident’ and ‘breeding season’), while removing polygons coded with other presence (e.g, extinct), origin (e.g. introduced, vagrant) and season (e.g. non-breeding, passage) attributes. We calculated range size as the area of the filtered distribution map transformed in a Mollweide equal-area projection. In addition, four predictors were extracted from BirdLife International (2022): generation length, migratory status, forest dependency and habitat breadth (calculated as the number of major habitat types that are coded as suitable for each species).

Morphological traits relating to beak length (from tip to nares), body mass and hand-wing index were extracted from AVONET (Tobias et al., 2022), alongside ecological information on trophic niches (aggregated into four classes: herbivore, omnivore, invertivore, and carnivore; Table S1). Insularity and clutch size were obtained from Tobias and Pigot (2019), and information on the diurnal/nocturnal activity of birds was obtained from Wilman et al. (2014). To address taxonomic mismatches, we matched all taxa to the taxonomy used by BirdLife International, using the synonym table from Tobias et al., (2022). The remaining 200 taxonomic mismatches were then corrected manually using the synonyms documented in BirdLife International (2022). A table of these matches is included in the supplementary tables provided with this article.

### B. Extrinsic factors

Similarly to the range size calculation, in measuring extrinsic variables we only considered breeding range for consistency among migratory and resident species. We used a raster layer of percentage tree-canopy cover in 2018 and changes in cover during 2000-2018 at 300-m resolution from Remelgado and Meyer (Under review; using Landsat data to correct some biases in global forest cover maps). We extracted the median value of these predictors within the range of each species. Similarly, we calculated the median value of range-wide cropland coverage (in 2019) and cropland changes from 2003 to 2019 obtained from Potapov et al. (2022).

We also calculated the median human population density within each species’ range, and the difference between median density in 2015 *vs* 2000, using data from the Global Human Settlement product (GHS; Schiavina et al., 2019). To account for direct pressures that species can face in the rural environment (as defined by the GHS product), we also calculated the proportion of the human population living in rural areas within each species’ range (Schiavina et al., 2019). As human accessibility can also determine the level of disturbance to which species are exposed, we extracted data from the global map of travel time to cities (Weiss et al., 2018) and calculated the median value of pixels contained within each species’ distribution. Finally, we extracted the countries of occurrence of each species from BirdLife International (2022) and calculated the resulting median Gross Domestic Product (GDP) per capita from WorldBank data (Worldbank, 2021) as an index of human population development.

In order to account for climatic correlates of risk, we extracted the current value and difference between past and current values for two variables from the CHELSAcruts database version 1.0 (Karger et al., 2017; Karger and Zimmermann, 2018), choosing Mean annual temperature and Annual precipitation amount for their relevance in influencing species’ distributions and their ease of interpretation (Supplementary Methods Table S1). Using data from another study predicting Red List categories (Lucas et al. 2023), we calculated the average value of both bioclimatic variables over two periods based on CHELSA data (Karger et al. 2017): 1965-1995, to represent the past climate, and 2005-2014, to represent the current climate. We then extracted the median value of each variable within the species’ range at both periods. We finally calculated the difference between both time periods, as a proxy of recent climate change experienced by the species within their geographic range.

We extracted raster values within species’ distribution polygons using the R package ‘*exactextractr’* (Baston et al., 2022). Polygons were reprojected according to each raster’s original coordinate system before extraction in order to minimise raster reprojection. Variables that followed a skewed distribution were log-transformed and all numeric variables were scaled (Table S1).

We removed 2,206 species for which we could not extract and calculate all predictors (mainly due to gaps for clutch size and insularity (1,425 species missing data) and diurnality (1,205 species missing data), leaving a final dataset composed of 8,999 species.

### 2. IUCN Red List framework

We extracted from BirdLife International (2022) the Red List category assigned to each species under each criterion (see below), as well as the final listed category (Fig. 1; see Fig. S1 for spatial distribution of these criteria). For birds, generally all Red List criteria have been evaluated for all species (with the exception of criterion E which is excluded from our study). For threatened bird species (i.e. those qualifying as Critically Endangered, Endangered and Vulnerable) all criteria that qualify a species as threatened should be reported in BirdLife International (2022), not just the one resulting in the highest risk category, although this may be omitted in some specific cases. However, this is not the case for species qualifying as Near Threatened (e.g., data are not available on whether a species classified as Endangered under B1 qualifies as Near Threatened under B2). To account for this, we followed two approaches.

**Figure 1.**
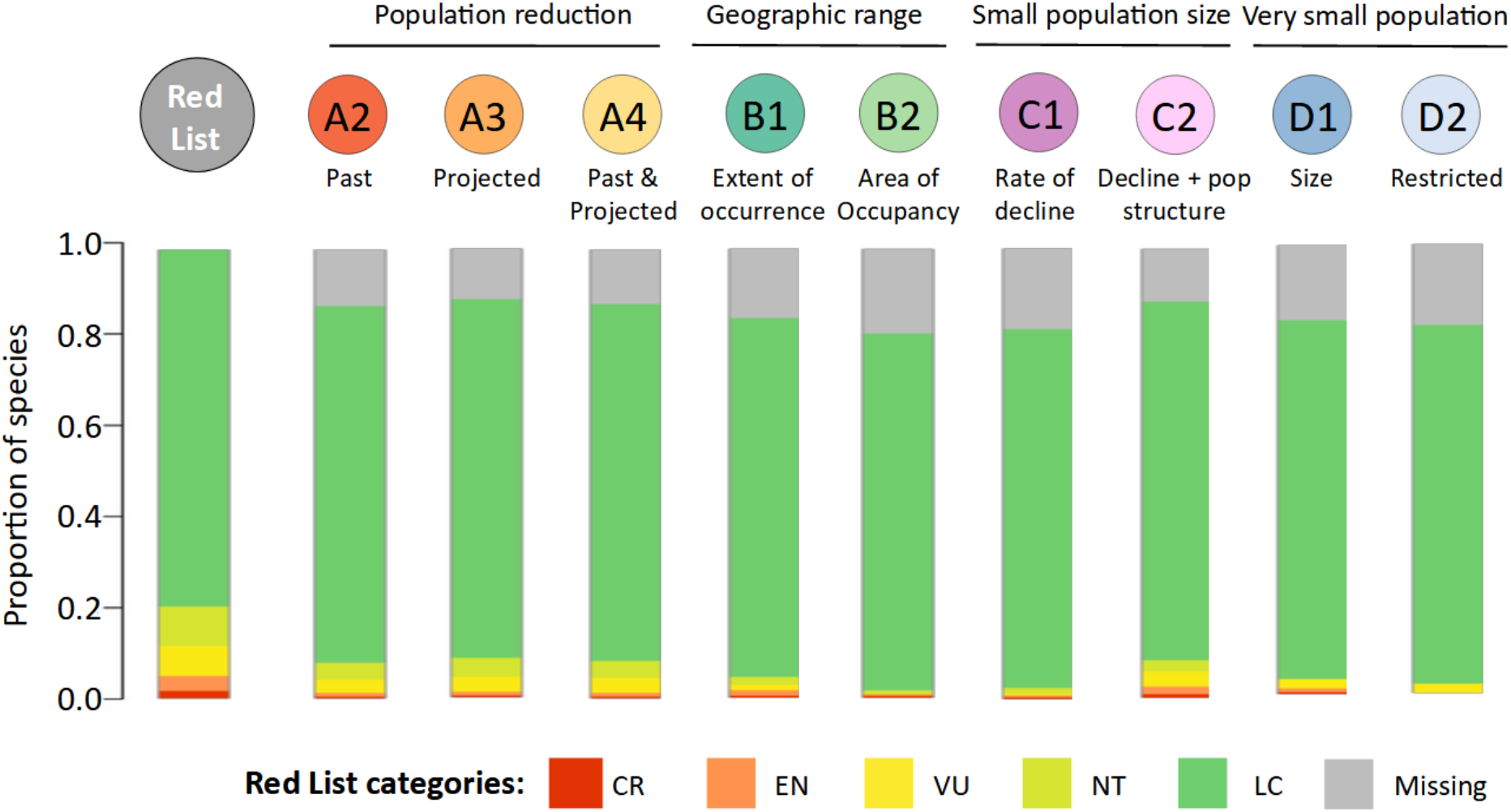
Proportion of the 8,999 bird species included in our analysis currently qualifying in each Red List category for each criterion. CR = Critically Endangered; EN = Endangered; VU= Vulnerable; NT = Near Threatened; LC = Least Concern. “Missing” applies to species qualifying as threatened for which the given criterion is not explicitly listed. Number of species per criterion is given in Table S2.

In the first, we classified a criterion as ‘missing data’ if not explicitly listed; the results presented in the main text, Fig.1 and Table S2 correspond to that assumption. In the second, we assumed that the species was Least Concern under a criterion unless it was explicitly listed; these results are reported in Table S3.

Only three species qualified as threatened under A1, hence the criterion has been excluded from the analysis.

### 3. Extinction risk modelling

We developed a “criterion-specific” modelling approach, in which we fitted a separate model for each Red List criterion. Each criterion is thus considered independently, and the Red List category met under that criterion is contrasted with the same set of extinction predictors. For comparison, we also fitted a “criterion-blind” model, as typically done in comparative extinction risk analyses, using the single listed species’ Red List category as the response variable.

To investigate the relationship between species traits, extrinsic factors and extinction risk, we used cumulative link models (CLM, also known as ordinal regression models) from the R package *‘ordinal’* (Christensen, 2019), which allow preservation of the ordinal structure of the Red List categories (Lucas et al., 2019, 2023; Luiz et al., 2016). We therefore considered the Red List category as an ordered categorical factor (LC < NT < VU < EN < CR; excluding all species with categories EX, EW, DD) and used it as the response variable and contrasted this with the predictors. Models varied from n = 7,278 for criterion B2 to n = 7,923 for criterion A3, with a total of 8,999 species included in the analysis. We checked that predictors were not highly correlated (Pearson correlation or Kruskal-gamma coefficients >|0.70|; Fig. S2). To adjust for unbalanced data (Fig. 1), we calculated the proportion of threatened and not threatened species under each criterion, and weighted our models by the proportion of species in the opposite category (i.e., species with categories VU, EN, or CR were weighted by the proportion of not threatened species and species with categories LC, or NT were weighted by the proportion of threatened species). A backward stepwise model selection was performed using the *step* function from the R package ‘*stats’* (R Core Team, 2020), in order to find the subset of variables that minimise the Akaike Information Criterion for each criteria. The proportional odds assumption of a linear relationship was not always met, but this should not impact our results substantially (see Supplementary Methods S2).

As the number of species listed in EN and CR categories was very small for some criteria (Fig. 1), we anticipated predicting specific categories could be challenging (Table S4); thus, for validation we focused on a simplified prediction: whether a species was classified as threatened or not. We used a method of taxonomic block validation in which we iteratively excluded one taxonomic family from the data used to train the model, and then used the model to predict the Red List binary threat level of the species in the left-out family. A species was predicted as threatened under a given criterion if the sum of the probability to be CR, EN, and VU was >0.5, and predicted as not threatened otherwise. We then compared the predictions with the actual Red List categorization under that specific criterion (assuming that the current Red List category of each species is correct for each criterion). Specificity and sensitivity were defined respectively as the percentage of not threatened species (LC, NT) correctly classified as such, and the percentage of threatened species (VU, EN, CR) correctly classified as such. Following Red List guidelines (IUCN Standards and Petitions Committee, 2022), we assigned a “combined” category to each species based on the nine criterion-specific models as the highest category predicted among models; consequently, a species was classified as not threatened only it was not predicted as threatened in any of the criteria models. We also report the performance of our models at predicting the specific Red List category for each species (assigning to each species the category with highest probability; Table S4).

To measure and compare the overall performance of both modelling approaches, we used the True Skill Statistic (TSS), defined as: Sensitivity + Specificity - 1. TSS takes into account both omission and commission errors, and ranges from −1 to +1, where +1 indicates perfect agreement, values greater than 0.5 indicate a good performance and values of 0 (or less) indicate a performance no better (or worse) than random (Allouche et al., 2006).

## Results

### 1. Model performance

Models performance in predicting extinction risk greatly varies among criteria (Fig. 2). The model predicting criterion B1, related to the extent of occurrence, had the highest TSS score (0.87), followed by criteria D2, D1, B2, C1, C2 (0.69 to 0.79). All these models had TSS scores higher than the criterion-blind approach (0.58), meaning that models are better at predicting extinction risk for single criteria than for the overall categories. For most criteria, these high TSS scores were the result of both higher specificity and higher sensitivity (Fig. 2). Conversely, models predicting criteria A2-4, related to population declines, showed the lowest TSS scores among all criteria (0.50-0.57), and they were slightly lower than the criterion-blind model.

**Figure 2.**
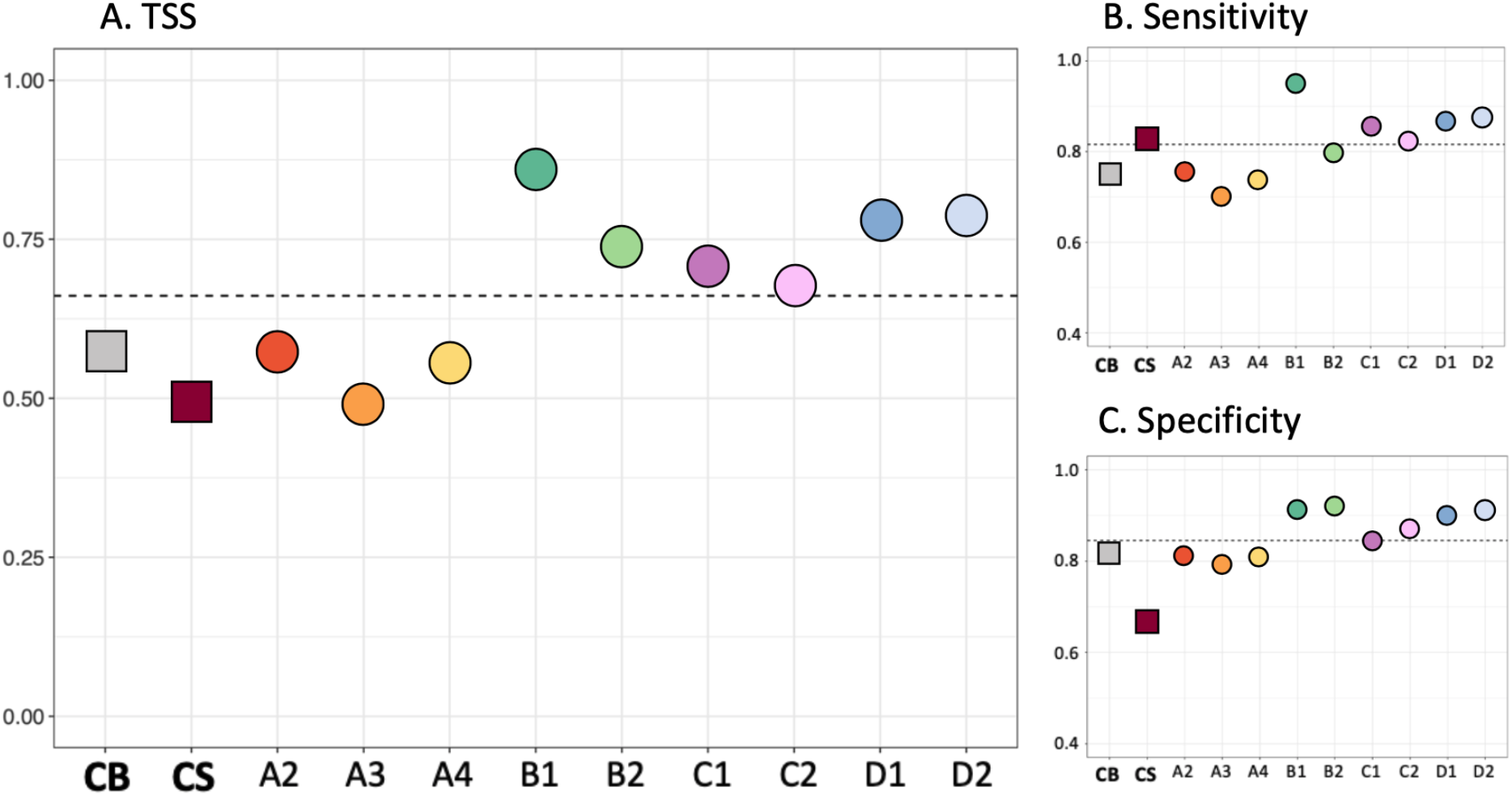
Comparison of model performances. The left-hand side of each plot compares the performance of the combined criterion-specific models (referred to as “**CS**”) with that of the criterion-blind approach to comparative extinction risk modelling (referred to as “**CB**”), while the right-hand side presents the performance of each criterion-specific model. A. True Skill Statistic [-1,1]; B. Sensitivity [0,1], proportion of threatened species correctly classified; C. Specificity [0,1], proportion of not threatened species correctly classified. Dotted lines represent the mean value obtained from the nine independent criteria-specific models.

Following Red List guidelines, we assigned a “combined” category to each species, as the highest extinction risk category from any of the nine individual criterion-specific models, and found that this substantially reduced the TSS (0.50) compared with applying the models individually for each criterion (average model TSS = 0.69; Figure 2A). This is largely due to lower specificity compared to the criterion-blind approach (0.69 vs. 0.82 probability of correctly classifying a not threatened species. Fig. 2C; Table S2), this is partly explained because a species had to be classified as not threatened under each of the nine applied criterion-explicit models in order to fall in this group overall. In contrast, using a criterion-specific approach resulted in 0.83 probability of correctly classifying threatened species, compared with 0.76 for the criterion-blind approach (Figure 2B; Table S2).

Considering “missing” criterion-specific categories as LC (Table S3) or predicting at the category level rather than binarily contrasting threatened vs not-threatened (Table S4) both resulted in lower performances.

### 2. Drivers of extinction risk

The criterion-blind approach showed positive relationships between extinction risk and body mass, carnivore trophic niche, high forest dependency and GDP per capita, while showing negative relationships with insularity, range size or percentage canopy cover (Figure 3). Some of these relationships were also detected by criteria-specific models. For instance, carnivorous species were generally associated with higher levels of extinction risk, while species with larger range size or with distributions that had a high average travel time to cities were less at risk on average.

**Figure 3.**
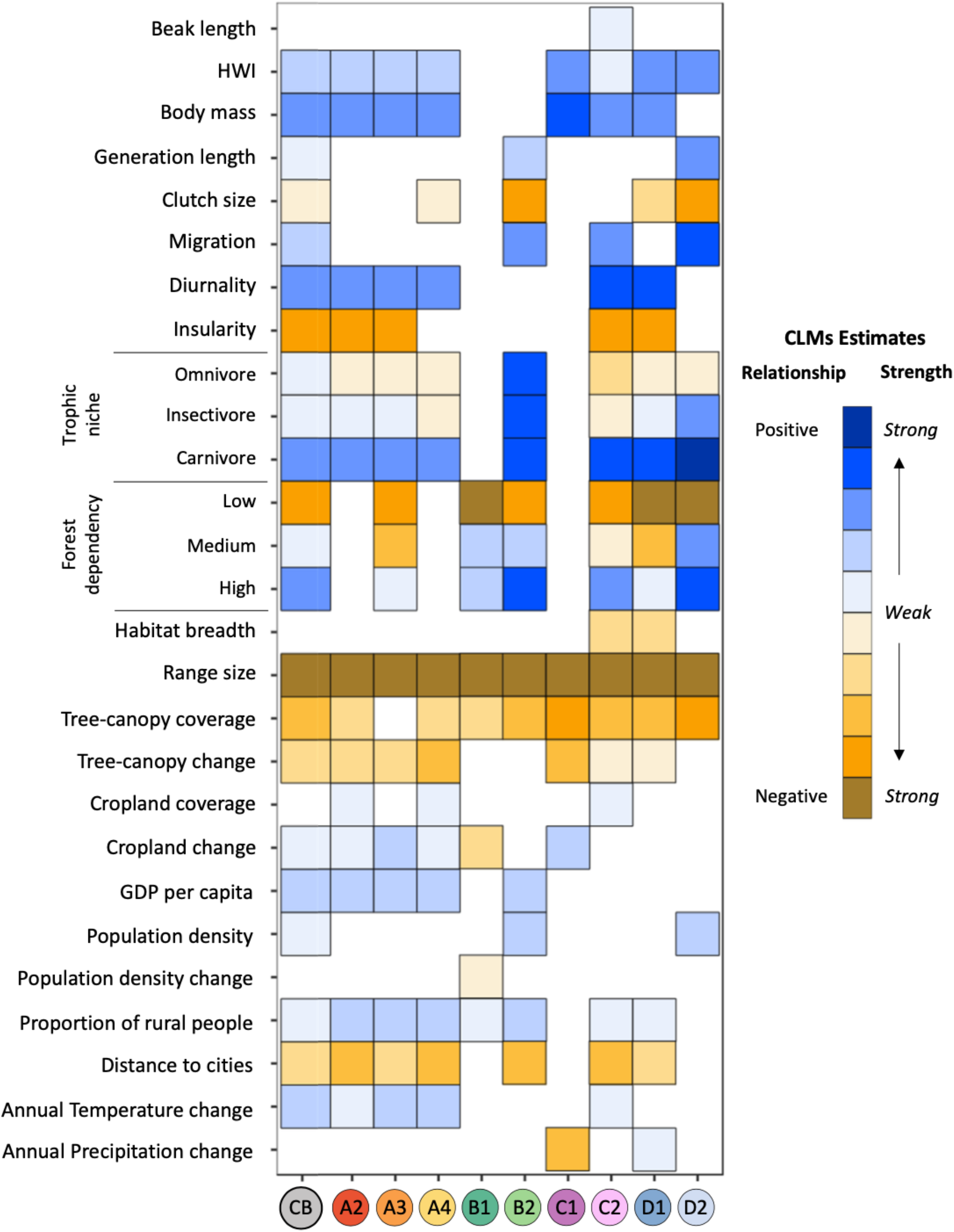
Heatmap of predictor importance associated with extinction risk under each criterion. Rows indicate predictors of extinction risk and columns relate to criterion-specific models. The ‘CB’ model describes the criterion-blind approach of extinction risk modelling. Colour indicates the sign (orange for negative and blue for positive) with darker tones indicating stronger relationships. Both positive and negative values were divided into 5 equal groups according to the intervals: -3.55, -0.957, -0.529, -0.291, -0.18, 0 for negative estimates and 0, 0.182, 0.387, 1.16, 3.02, 12.2 for positive estimates (blanks indicate that the predictor has not been retained in the optimal model after predictor selection; see Methods section 3).

In contrast, the importance and significance of other predictors were idiosyncratic between criteria. For instance, body mass generally correlated positively with extinction risk for criteria related to rapid population declines or small population size (A2, A3, A4, C1, C2, D1) while body mass had no influence on criteria related to restricted geographic range (criteria B1, B2, D2). Conversely, high forest dependency was associated with increased extinction risk for criteria B2, C2 and D2, all relating to small population size, restricted area of occupancy and subpopulation structure, but did not influence criteria based on rates of decline alone. Extrinsic factors were mainly important to predict criteria related to population reductions (A2, A3, A4), for instance change in annual temperature correlating positively with extinction risk for criteria A2-4 and C2, and GDP per capita correlating positively with criteria A2-4 and B2.

### 3. Criterion-specific approach to prioritise re-assessments

The criterion-blind model predicts that 18% of species currently classified as not threatened (N=1,408) may be threatened (Table 1), but this percentage almost doubled (to 33%, N=2,644) under the criterion-specific model. Conversely, we predicted 114 threatened species as not threatened (233 with the criterion-blind model). Predictions for all models and all species are provided in Supplementary Information.

**Table 1.** Red List category prediction of criterion-blind and criterion-specific models compared with the current binarized category (Threatened for Vulnerable, Endangered, Critically Endangered; Not threatened for Least Concern, Near Threatened). The prediction for the criterion-specific method corresponds to the prediction after combining results from the nine individual criterion models.

| IUCN Red List | Prediction criterion-blind |  | Prediction criterion-specific |  |
| --- | --- | --- | --- | --- |
|  | Not threatened | Threatened | Not threatened | Threatened |
| Not threatened | 6,531 (72%) | 1,408 (16%) | 5,295 (59%) | 2,644 (29%) |
| Threatened | 233 (3%) | 827 (9%) | 114 (1%) | 946 (11%) |

## Discussion

In this study, we developed a modelling approach that partitions extinction risk according to individual Red List criteria and compared it with a criterion-blind approach. On average, modelling individual criteria performed better than the criterion-blind approach, with higher performance for six criteria (especially criterion B1), while three provided similar or marginally lower performance (criteria A2-A4). This result highlights that predicting extinction risk under some criteria may be intrinsically difficult, at least using the predictors considered here. In particular, criteria related to population trends (especially A3 related to future trends) are more difficult to predict. With these models, we can enhance our understanding of the mechanisms underlying observed correlations and ultimately, point to distinct drivers of risk.

Combining the nine criteria-specific models to obtain a single prediction per species led to substantially greater sensitivity (i.e. more likely prediction of of threatened species as threatened, Figure 2B) but lower specificity than the classical criterion-blind approach (see, for example, White-winged Petrel *Pterodroma leucoptera* in Figure 4A). Because one of the primary goals of automated extinction risk methods is to identify species likely to be at risk of extinction (but not currently recognised as such) to prioritise their reassessment (Cardillo and Meijaard, 2012), a model with high sensitivity will be more valuable than a model with similar TSS higher specificity (Cazalis et al. 2022). Previous extinction risk models have typically predicted threatened species less accurately than not threatened species (Murray et al., 2014; Di Marco, 2022). Our results show that a reason behind this observation may be the omission of the diversity of reasons why a species is considered threatened in the Red List, which is represented by the multiple Red List criteria. However, combining nine criterion-specific models decreased the overall specificity of the prediction (see, for example, Sangihe Scops-owl *Otus collari* in Figure 4B), resulting in an overestimation of the proportion of threatened species, and a slightly lower TSS in comparison with the criterion-blind approach (Figure 2A). This result is explained by the fact that, following Red List guidelines, a species was classified as not threatened only if predicted as such by all nine criterion-specific models. This is well aligned with the Red List process and a precautionary approach, but makes our approach sensitive to misclassification. Hence, increasing the specificity of criterion-specific models is a priority for the future. Possible ways of achieving this include improving the performance of the individual models with additional covariates (for example, relating to hunting/trapping pressure for criteria A2-4), or developing an approach to combine individual models that accumulates fewer errors from individual models that misclassify a species as threatened under a given criterion. In time, such improvements may also enable predictions at the category level (while we limit here to a binary threatened/not threatened prediction).

**Figure 4.**
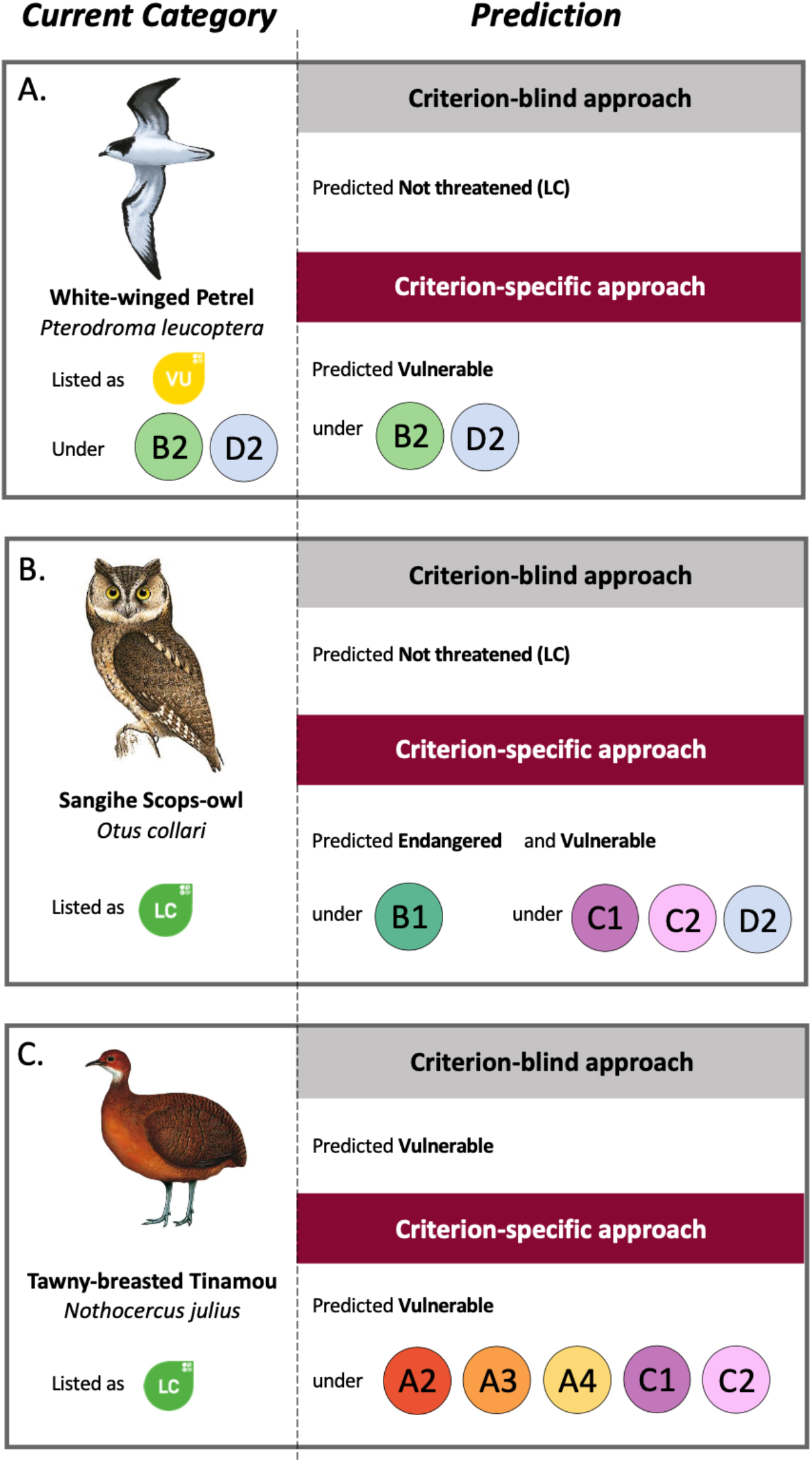
Comparison of outputs for selected species from a criterion-blind approach and a criterion-specific approach to comparative extinction risk analysis. The three panels show different species, with their current categories in the Red List. Predictions are distinguished according to the approaches, indicating the categories predicted by the models and the criteria triggered for the criterion-specific approach. Illustration by A. Juan Varela, B Ian Willis. C. Lluís Sanz, source: © 2022 Cornell University

In accordance with previous studies, our criterion-blind model identified extinction risk as correlating positively with e.g., body mass, generation length, high forest dependency, and tree-canopy change, and negatively with e.g., clutch size, range size, and distance to cities (Gaston and Blackburn, 1995; Olah et al., 2018; Richards et al., 2021; Ripple et al., 2017; Tobias and Pigot, 2019; White and Bennett, 2015). Our findings highlight the importance of considering separately the multiple processes underlying patterns of extinction (Figs 3, S1). They reveal that increases in annual temperature across species’ ranges are of particular importance for criteria related to population decline (A2-A4, and to some extent C2), consistent with the predicted role of climate change in driving declines in abundance and increases in species’ extinction risk (Şekercioğlu et al., 2012). Additionally, body mass is of particular importance for almost all criteria except those related to geographic range size (B1 and B2), likely because larger species typically require larger range sizes to retain viable populations (Carvajal-Quintero et al. 2023). The diversity of relationships between predictors and criteria, along with their ecological meaning, emphasises the importance of accounting for heterogeneity in the predictability of Red List criteria, rather than considering them as equally predictable as assumed in criterion-blind comparative extinction risk analyses. They also highlight that our criterion-specific approach can help better understand the diverse mechanisms associated with extinction risk. Further, a better approximation of the causal relationships underlying species classification under different Red List categories can improve our ability to forecast status change based on changes in the drivers.

By modelling criteria separately, we increase the applicability of comparative extinction risk models (Cazalis et al., 2022; Owens and Bennett, 2000; Ripple et al., 2017). Red List assessors are required to assess each species against all criteria for which there is sufficient information (IUCN Standards and Petitions Committee, 2022). Therefore, while our model outputs do not fundamentally change the red-listing process, they allow for various sources of information to be taken into account by assessors, thus, helping prioritise efforts to assess/reassess species identified as possibly being at risk, and identify knowledge gaps or opportunities for future research. For instance, the Least Concern Tawny-breasted Tinamou *Nothocercus julius* (Figure 4C) is predicted to be threatened (VU) by both the criterion-blind and the criterion-specific approaches. However, while the criterion-blind model offers no additional insight, our criterion-specific approach provides assessors with relevant information about why this species might be VU – namely that it might meet criteria A2, A3, A4, C1 and C2. Assessors could use this information to focus efforts on investigating past and future population trends to assess criteria A2-4 and population size and structure to assess criteria C1 and C2, which could be complemented with the specific contributions of different covariates that led to this prediction. If these data are not available, determining values for these parameters may be considered a priority for fieldwork and research.

Comparative extinction risk models have often been promoted as useful tools to provide a first prediction of extinction risk for species not yet included in the Red List (Darrah et al., 2017; Zizka et al., 2021a; Zizka et al., 2022), for Data Deficient species (Bland and Böhm, 2016; Borgelt et al., 2022; He et al., 2021; Luiz et al., 2016), or to prioritise reassessments (Lucas et al., 2023; Di Marco et al., 2014). But so far these have remained largely unmet promises, with hardly any uptake of these modelling approaches in the Red List (Cazalis et al. 2022). By informing the assessors about the specific criteria under which a species is likely to qualify, criterion-specific models could help accelerate the rate of Red List assessments, guide data collection efforts, and facilitate the growth and update of the Red List so that it can best inform conservation policies. We believe this is a promising avenue to reduce the historical research-implementation gap between comparative extinction risk model and the Red List assessment process.

## Supporting information

Birds taxonomic crosswalks

Criterion specific predictions

## Bibliography

Allouche, O., Tsoar, A., Kadmon, R., 2006. Assessing the accuracy of species distribution models: prevalence, kappa and the true skill statistic (TSS). Journal of Applied Ecology 43, 1223–1232. https://doi.org/10.1111/j.1365-2664.2006.01214.x

Bachman, S.P., Field, R., Reader, T., Raimondo, D., Donaldson, J., Schatz, G.E., Lughadha, E.N., 2019. Progress, challenges and opportunities for Red Listing. Biological Conservation 234, 45–55. https://doi.org/10.1016/j.biocon.2019.03.002

Baston, ISciences, LLC, 2022. exactextractr, Version:0.8.2.

Betts, J., Young, R.P., Hilton-Taylor, C., Hoffmann, M., Rodríguez, J.P., Stuart, S.N., Milner-Gulland, E.J., 2020. A framework for evaluating the impact of the IUCN Red List of threatened species. Conservation Biology 34, 632–643. https://doi.org/10.1111/cobi.13454

BirdLife International, 2022. IUCN Red List for birds. Downloaded from http://www.birdlife.orgon 15/06/2022.

BirdLife International and Handbook of the Birds of the World, 2021. Bird species distribution maps of the world. Version 2021.1. Available at http://datazone.birdlife.org/species/requestdis.

Bland, L.M., Böhm, M., 2016. Overcoming data deficiency in reptiles. Biological Conservation, Advancing reptile conservation: Addressing knowledge gaps and mitigating key drivers of extinction risk 204, 16–22. https://doi.org/10.1016/j.biocon.2016.05.018

Borgelt, J., Dorber, M., Høiberg, M.A., Verones, F., 2022. More than half of data deficient species predicted to be threatened by extinction. Commun Biol 5, 1–9. https://doi.org/10.1038/s42003-022-03638-9

Buchanan, G. M., Butchart, S. H. M., Dutson, G., Pilgrim, J. D., Steininger, M. K., Bishop, K. D., & Mayaux, P. (2008). Using remote sensing to inform conservation status assessment : Estimates of recent deforestation rates on New Britain and the impacts upon endemic birds. iological Conservation, 141(1), Article 1. https://doi.org/10.1016/j.biocon.2007.08.023

Cardillo, M., Meijaard, E., 2012. Are comparative studies of extinction risk useful for conservation? Trends in Ecology & Evolution 27, 167–171. https://doi.org/10.1016/j.tree.2011.09.013

Carvajal-Quintero, J., Comte, L., Giam, X., Olden, J. D., Brose, U., Erős, T., Filipe, A. F., Fortin, M., Irving, K., Jacquet, C., Larsen, S., Ruhi, A., Sharma, S., Villalobos, F., Tedesco, P. A., 2023. Scale of population synchrony confirms macroecological estimates of minimum viable range size. Ecology Letters, 26(2), 291–301. https://doi.org/10.1111/ele.14152

Cazalis, V., Di Marco, M., Butchart, S.H.M., Akçakaya, H.R., González-Suárez, M., Meyer, C., Clausnitzer, V., Böhm, M., Zizka, A., Cardoso, P., Schipper, A.M., Bachman, S.P., Young, B.E., Hoffmann, M., Benítez-López, A., Lucas, P.M., Pettorelli, N., Patoine, G., Pacifici, M., Jörger-Hickfang, T., Brooks, T.M., Rondinini, C., Hill, S.L.L., Visconti, P., Santini, L., 2022. Bridging the research-implementation gap in IUCN Red List assessments. Trends in Ecology & Evolution. https://doi.org/10.1016/j.tree.2021.12.002

Cazalis, V. (2022). Species richness response to human pressure hides important assemblage transformations. Proceedings of the National Academy of Sciences, 119(19), Article 19. https://doi.org/10.1073/pnas.2107361119

Cazalis, V., Santini, L., Lucas, P.M., González-Suárez, M., Hoffmann, M., Benítez-López, A., Pacifici, M., Schipper, A.M., Böhm, M., Zizka, A., Clausnitzer, V., Meyer, C., Jung, M., Butchart, S.H.M., Cardoso, P., Mancini, G., Akçakaya, H.R., Young, B.E., Patoine, G., Di Marco, M. (2023). Prioritizing the reassessment of Data Deficient species in the Red List. Accepted in Conservation Biology.

Chichorro, F., Juslén, A., Cardoso, P., 2019. A review of the relation between species traits and extinction risk. Biological Conservation 237, 220–229. https://doi.org/10.1016/j.biocon.2019.07.001

Chichorro, F., Correia, L., & Cardoso, P. (2022). Biological traits interact with human threats to drive extinctions : A modelling study. Ecological Informatics, 69, 101604. https://doi.org/10.1016/j.ecoinf.2022.101604

Christensen, R., 2019. “ordinal—Regression Models for Ordinal Data.” R package version 2019.12-10.

Damuth, J., 1981. Population density and body size in mammals. Nature 290, 699–700. https://doi.org/10.1038/290699a0

Darrah, S.E., Bland, L.M., Bachman, S.P., Clubbe, C.P., Trias-Blasi, A., 2017. Using coarse-scale species distribution data to predict extinction risk in plants. Diversity and Distributions 23, 435–447. https://doi.org/10.1111/ddi.12532

Davidson, A.D., Hamilton, M.J., Boyer, A.G., Brown, J.H., Ceballos, G., 2009. Multiple ecological pathways to extinction in mammals. Proceedings of the National Academy of Sciences 106, 10702–10705. https://doi.org/10.1073/pnas.0901956106

Di Marco, M., 2022. Reptile research shows new avenues and old challenges for extinction risk modelling. PLOS Biology 20, e3001719. https://doi.org/10.1371/journal.pbio.3001719

Di Marco, M., Buchanan, G.M., Szantoi, Z., Holmgren, M., Grottolo Marasini, G., Gross, D., Tranquilli, S., Boitani, L., Rondinini, C., 2014. Drivers of extinction risk in African mammals: the interplay of distribution state, human pressure, conservation response and species biology. Phil. Trans. R. Soc. B 369, 20130198. https://doi.org/10.1098/rstb.2013.0198

Gaston, K.J., Blackburn, T.M., 1995. Birds, Body Size and the Threat of Extinction. Philosophical Transactions: Biological Sciences 347, 205–212.

González-Suárez, M., Gómez, A., Revilla, E., 2013. Which intrinsic traits predict vulnerability to extinction depends on the actual threatening processes. Ecosphere 4, art76. https://doi.org/10.1890/ES12-00380.1

He, F., Langhans, S.D., Zarfl, C., Wanke, R., Tockner, K., Jähnig, S.C., 2021. Combined effects of life-history traits and human impact on extinction risk of freshwater megafauna. Conservation Biology 35, 643–653. https://doi.org/10.1111/cobi.13590

Hoffmann, M., Hilton-Taylor, C., Angulo, A., Böhm, M., Brooks, T.M., Butchart, S.H.M., Carpenter, K.E., Chanson, J., Collen, B., Cox, N.A., Darwall, W.R.T., Dulvy, N.K., Harrison, L.R., Katariya, V., Pollock, C.M., Quader, S., Richman, N.I., Rodrigues, A.S.L., Tognelli, M.F., Vié, J.-C., Aguiar, J.M., Allen, D.J., Allen, G.R., Amori, G., Ananjeva, N.B., Andreone, F., Andrew, P., Ortiz, A.L.A., Baillie, J.E.M., Baldi, R., Bell, B.D., Biju, S.D., Bird, J.P., Black-Decima, P., Blanc, J.J., Bolaños, F., Bolivar-G., W., Burfield, I.J., Burton, J.A., Capper, D.R., Castro, F., Catullo, G., Cavanagh, R.D., Channing, A., Chao, N.L., Chenery, A.M., Chiozza, F., Clausnitzer, V., Collar, N.J., Collett, L.C., Collette, B.B., Fernandez, C.F.C., Craig, M.T., Crosby, M.J., Cumberlidge, N., Cuttelod, A., Derocher, A.E., Diesmos, A.C., Donaldson, J.S., Duckworth, J.W., Dutson, G., Dutta, S.K., Emslie, R.H., Farjon, A., Fowler, S., Freyhof, J., Garshelis, D.L., Gerlach, J., Gower, D.J., Grant, T.D., Hammerson, G.A., Harris, R.B., Heaney, L.R., Hedges, S.B., Hero, J.-M., Hughes, B., Hussain, S.A., Icochea M. J., Inger, R.F., Ishii, N., Iskandar, D.T., Jenkins, R.K.B., Kaneko, Y., Kottelat, M., Kovacs, K.M., Kuzmin, S.L., La Marca, E., Lamoreux, J.F., Lau, M.W.N., Lavilla, E.O., Leus, K., Lewison, R.L., Lichtenstein, G., Livingstone, S.R., Lukoschek, V., Mallon, D.P., McGowan, P.J.K., McIvor, A., Moehlman, P.D., Molur, S., Alonso, A.M., Musick, J.A., Nowell, K., Nussbaum, R.A., Olech, W., Orlov, N.L., Papenfuss, T.J., Parra-Olea, G., Perrin, W.F., Polidoro, B.A., Pourkazemi, M., Racey, P.A., Ragle, J.S., Ram, M., Rathbun, G., Reynolds, R.P., Rhodin, A.G.J., Richards, S.J., Rodríguez, L.O., Ron, S.R., Rondinini, C., Rylands, A.B., Sadovy de Mitcheson, Y., Sanciangco, J.C., Sanders, K.L., Santos-Barrera, G., Schipper, J., Self-Sullivan, C., Shi, Y., Shoemaker, A., Short, F.T., Sillero-Zubiri, C., Silvano, D.L., Smith, K.G., Smith, A.T., Snoeks, J., Stattersfield, A.J., Symes, A.J., Taber, A.B., Talukdar, B.K., Temple, H.J., Timmins, R., Tobias, J.A., Tsytsulina, K., Tweddle, D., Ubeda, C., Valenti, S.V., Paul van Dijk, P., Veiga, L.M., Veloso, A., Wege, D.C., Wilkinson, M., Williamson, E.A., Xie, F., Young, B.E., Akçakaya, H.R., Bennun, L., Blackburn, T.M., Boitani, L., Dublin, H.T., da Fonseca, G.A.B., Gascon, C., Lacher, T.E., Mace, G.M., Mainka, S.A., McNeely, J.A., Mittermeier, R.A., Reid, G.M., Rodriguez, J.P., Rosenberg, A.A., Samways, M.J., Smart, J., Stein, B.A., Stuart, S.N., 2010. The Impact of Conservation on the Status of the World’s Vertebrates. Science 330, 1503–1509. https://doi.org/10.1126/science.1194442

IUCN, 2022. The IUCN Red List of Threatened Species. Version 2022-2.

IUCN, 2012. IUCN Red List Categories and Criteria: Version 3.1. Second edition (Gland, Switzeland and Cambridge, UK: IUCN).

IUCN Standards and Petitions Committee. (2022). Guidelines for Using the IUCN Red List Categories and Criteria. Version 15.1. Prepared by the Standards and Petitions Committee. Downloadable from http://www.iucnredlist.org/documents/RedListGuidelines.pdf.

Jones, M.J., Fielding, A., Sullivan, M., 2006. Analysing Extinction Risk in Parrots using Decision Trees. Biodivers Conserv 15, 1993–2007. https://doi.org/10.1007/s10531-005-4316-1

Juffe-Bignoli, D., Brooks, T.M., Butchart, S.H.M., Jenkins, R.B., Boe, K., Hoffmann, M., Angulo, A., Bachman, S., Böhm, M., Brummitt, N., Carpenter, K.E., Comer, P.J., Cox, N., Cuttelod, A., Darwall, W.R.T., Marco, M.D., Fishpool, L.D.C., Goettsch, B., Heath, M., Hilton-Taylor, C., Hutton, J., Johnson, T., Joolia, A., Keith, D.A., Langhammer, P.F., Luedtke, J., Lughadha, E.N., Lutz, M., May, I., Miller, R.M., Oliveira-Miranda, M.A., Parr, M., Pollock, C.M., Ralph, G., Rodríguez, J.P., Rondinini, C., Smart, J., Stuart, S., Symes, A., Tordoff, A.W., Woodley, S., Young, B., Kingston, N., 2016. Assessing the Cost of Global Biodiversity and Conservation Knowledge. PLOS ONE 11, e0160640. https://doi.org/10.1371/journal.pone.0160640

Karger, D.N., Conrad, O., Böhner, J., Kawohl, T., Kreft, H., Soria-Auza, R.W., Zimmermann, N.E., Linder, H.P., Kessler, M., 2017. Climatologies at high resolution for the earth’s land surface areas. Sci Data 4, 170122. https://doi.org/10.1038/sdata.2017.122

Karger, D.N., Zimmermann, N.E., 2018. CHELSAcruts - High resolution temperature and precipitation timeseries for the 20th century and beyond. https://doi.org/10.16904/ENVIDAT.159

Kassambara, A., Mundt, F., 2020. factoextra: Extract and Visualize the Results of Multivariate Data Analyses. R package version 1.0.7.

Lees, A. C., Haskell, L., Allinson, T., Bezeng, S. B., Burfield, I. J., Renjifo, L. M., Rosenberg, K. V., Viswanathan, A., & Butchart, S. H. M. (2022). State of the World’s Birds. Annual Review of Environment and Resources, 47(1), Article 1. https://doi.org/10.1146/annurev-environ-112420-014642

Lucas, P.M., Di Marco, M., Cazalis, V., Luedtke, J., Brown, M., Langhammer, P., Neam, K., Mancini, G., Santini, L., 2023. Testing the predictive performance of comparative extinction risk models to support the global amphibian assessment. BioRxiv. https://doi.org/10.1101/2023.02.08.526823

Lucas, P.M., González-Suárez, M., Revilla, E., 2019. Range area matters, and so does spatial configuration: predicting conservation status in vertebrates. Ecography 42, 1103–1114. https://doi.org/10.1111/ecog.03865

Luiz, O.J., Woods, R.M., Madin, E.M.P., Madin, J.S., 2016. Predicting IUCN Extinction Risk Categories for the World’s Data Deficient Groupers (Teleostei: Epinephelidae). Conservation Letters 9, 342–350. https://doi.org/10.1111/conl.12230

Mace, G.M., Collar, N.J., Gaston, K.J., Hilton-Taylor, C., Akçakaya, H.R., Leader-Williams, N., Milner-Gulland, E.J., Stuart, S.N., 2008. Quantification of Extinction Risk: IUCN’s System for Classifying Threatened Species. Conservation Biology 22, 1424–1442. https://doi.org/10.1111/j.1523-1739.2008.01044.x

Murray, K.A., Verde Arregoitia, L.D., Davidson, A., Di Marco, M., Di Fonzo, M.M.I., 2014. Threat to the point: improving the value of comparative extinction risk analysis for conservation action. Global Change Biology 20, 483–494. https://doi.org/10.1111/gcb.12366

Newsome, T.M., Wolf, C., Nimmo, D.G., Kopf, R.K., Ritchie, E.G., Smith, F.A., Ripple, W.J., 2020. Constraints on vertebrate range size predict extinction risk. Global Ecology and Biogeography 29, 76–86. https://doi.org/10.1111/geb.13009

Olah, G., Theuerkauf, J., Legault, A., Gula, R., Stein, J., Butchart, S., O’Brien, M., Heinsohn, R., 2018. Parrots of Oceania – a comparative study of extinction risk. Emu - Austral Ornithology 118, 94–112. https://doi.org/10.1080/01584197.2017.1410066

Owens, I.P.F., Bennett, P.M., 2000. Ecological basis of extinction risk in birds: Habitat loss versus human persecution and introduced predators. Proceedings of the National Academy of Sciences 97, 12144–12148. https://doi.org/10.1073/pnas.200223397

Potapov, P., Turubanova, S., Hansen, M.C., Tyukavina, A., Zalles, V., Khan, A., Song, X.-P., Pickens, A., Shen, Q., Cortez, J., 2022. Global maps of cropland extent and change show accelerated cropland expansion in the twenty-first century. Nat Food 3, 19–28. https://doi.org/10.1038/s43016-021-00429-z

Purvis, A., Gittleman, J. L., Cowlishaw, G., & Mace, G. M. (2000). Predicting extinction risk in declining species. Proceedings of the Royal Society B: Biological Sciences, 267(1456), Article 1456. https://doi.org/10.1098/rspb.2000.1234

R Core Team, 2020. R: A language and environment for statistical computing.

Remelgado, R., Meyer, C. (Under review). Global dynamics in tree-canopy density over three decades. Under review in Global Change Biology

Richards, C., Cooke, R.S.C., Bates, A.E., 2021. Biological traits of seabirds predict extinction risk and vulnerability to anthropogenic threats. Global Ecology and Biogeography 30, 973–986. https://doi.org/10.1111/geb.13279

Ripple, W.J., Wolf, C., Newsome, T.M., Hoffmann, M., Wirsing, A.J., McCauley, D.J., 2017. Extinction risk is most acute for the world’s largest and smallest vertebrates. Proceedings of the National Academy of Sciences 114, 10678–10683. https://doi.org/10.1073/pnas.1702078114

Rodrigues, A.S.L., Pilgrim, J.D., Lamoreux, J.F., Hoffmann, M., Brooks, T.M., 2006. The value of the IUCN Red List for conservation. Trends in Ecology & Evolution 21, 71–76. https://doi.org/10.1016/j.tree.2005.10.010

Rondinini, C., Di Marco, M., Visconti, P., Butchart, S.H.M., Boitani, L., 2014. Update or Outdate: Long-Term Viability of the IUCN Red List. Conservation Letters 7, 126–130. https://doi.org/10.1111/conl.12040

Santini, L., Butchart, S.H.M., Rondinini, C., Benítez-López, A., Hilbers, J.P., Schipper, A.M., Cengic, M., Tobias, J.A., Huijbregts, M.A.J., 2019. Applying habitat and population-density models to land-cover time series to inform IUCN Red List assessments. Conservation Biology 33, 1084–1093. https://doi.org/10.1111/cobi.13279

Santini, L., Benítez-López, A., Dormann, C. F., & Huijbregts, M. A. (2022). Population density estimates for terrestrial mammal species. Global Ecology and Biogeography, 31(5), 978–994.

Schiavina, M., Freire, S., MacManus, K., 2019. GHS-POP R2019A - GHS population grid multitemporal (1975-1990-2000-2015). https://doi.org/10.2905/0C6B9751-A71F-4062-830B-43C9F432370F

Şekercioğlu, Ç.H., Primack, R.B., Wormworth, J., 2012. The effects of climate change on tropical birds. Biological Conservation 148, 1–18. https://doi.org/10.1016/j.biocon.2011.10.019

Senn, S., Julious, S., 2009. Measurement in clinical trials: A neglected issue for statisticians? Statistics in Medicine 28, 3189–3209. https://doi.org/10.1002/sim.3603

Silva, M., & Downing, J. A. (1995). The allometric scaling of density and body mass: A non-linear relationship for terrestrial mammals.American Naturalist, 145, 704–727. https://doi.org/10.1086/285764

Tobias, J.A., Pigot, A.L., 2019. Integrating behaviour and ecology into global biodiversity conservation strategies. Philosophical Transactions of the Royal Society B: Biological Sciences 374, 20190012. https://doi.org/10.1098/rstb.2019.0012

Tobias, J.A., Sheard, C., Pigot, A.L., Devenish, A.J.M., Yang, J., Sayol, F., Neate-Clegg, M.H.C., Alioravainen, N., Weeks, T.L., Barber, R.A., Walkden, P.A., MacGregor, H.E.A., Jones, S.E.I., Vincent, C., Phillips, A.G., Marples, N.M., Montaño-Centellas, F.A., Leandro-Silva, V., Claramunt, S., Darski, B., Freeman, B.G., Bregman, T.P., Cooney, C.R., Hughes, E.C., Capp, E.J.R., Varley, Z.K., Friedman, N.R., Korntheuer, H., Corrales-Vargas, A., Trisos, C.H., Weeks, B.C., Hanz, D.M., Töpfer, T., Bravo, G.A., Remeš, V., Nowak, L., Carneiro, L.S., Moncada R. A.J., Matysioková, B., Baldassarre, D.T., Martínez-Salinas, A., Wolfe, J.D., Chapman, P.M., Daly, B.G., Sorensen, M.C., Neu, A., Ford, M.A., Mayhew, R.J., Fabio Silveira, L., Kelly, D.J., Annorbah, N.N.D., Pollock, H.S., Grabowska-Zhang, A.M., McEntee, J.P., Carlos T. Gonzalez, J., Meneses, C.G., Muñoz, M.C., Powell, L.L., Jamie, G.A., Matthews, T.J., Johnson, O., Brito, G.R.R., Zyskowski, K., Crates, R., Harvey, M.G., Jurado Zevallos, M., Hosner, P.A., Bradfer-Lawrence, T., Maley, J.M., Stiles, F.G., Lima, H.S., Provost, K.L., Chibesa, M., Mashao, M., Howard, J.T., Mlamba, E., Chua, M.A.H., Li, B., Gómez, M.I., García, N.C., Päckert, M., Fuchs, J., Ali, J.R., Derryberry, E.P., Carlson, M.L., Urriza, R.C., Brzeski, K.E., Prawiradilaga, D.M., Rayner, M.J., Miller, E.T., Bowie, R.C.K., Lafontaine, R.-M., Scofield, R.P., Lou, Y., Somarathna, L., Lepage, D., Illif, M., Neuschulz, E.L., Templin, M., Dehling, D.M., Cooper, J.C., Pauwels, O.S.G., Analuddin, K., Fjeldså, J., Seddon, N., Sweet, P.R., DeClerck, F.A.J., Naka, L.N., Brawn, J.D., Aleixo, A., Böhning-Gaese, K., Rahbek, C., Fritz, S.A., Thomas, G.H., Schleuning, M., 2022. AVONET: morphological, ecological and geographical data for all birds. Ecology Letters 25, 581–597. https://doi.org/10.1111/ele.13898

Weiss, D.J., Nelson, A., Gibson, H.S., Temperley, W., Peedell, S., Lieber, A., Hancher, M., Poyart, E., Belchior, S., Fullman, N., Mappin, B., Dalrymple, U., Rozier, J., Lucas, T.C.D., Howes, R.E., Tusting, L.S., Kang, S.Y., Cameron, E., Bisanzio, D., Battle, K.E., Bhatt, S., Gething, P.W., 2018. A global map of travel time to cities to assess inequalities in accessibility in 2015. Nature 553, 333–336. https://doi.org/10.1038/nature25181

White, R.L., Bennett, P.M., 2015. Elevational Distribution and Extinction Risk in Birds. PLOS ONE 10, e0121849. https://doi.org/10.1371/journal.pone.0121849

Williams, B.A., Watson, J.E.M., Butchart, S.H.M., Ward, M., Brooks, T.M., Butt, N., Bolam, F.C., Stuart, S.N., Mair, L., McGowan, P.J.K., Gregory, R., Hilton-Taylor, C., Mallon, D., Harrison, I., Simmonds, J.S., 2021. A robust goal is needed for species in the Post-2020 Global Biodiversity Framework. Conservation Letters 14, e12778. https://doi.org/10.1111/conl.12778

Wilman, H., Belmaker, J., Simpson, J., de la Rosa, C., Rivadeneira, M.M., Jetz, W., 2014. EltonTraits 1.0: Species-level foraging attributes of the world’s birds and mammals. Ecology 95, 2027–2027. https://doi.org/10.1890/13-1917.1

Worldbank, 2021. GDP per Capita. World Bank Development Indicators, The World Bank Group, accessed at https://data.worldbank.org/indicator/NY.GDP.CAP.CD 01-11-2021.

Zizka, A., Barratt, C.D., Ritter, C.D., Joerger-Hickfang, T., Zizka, V.M.A., 2021a. Existing approaches and future directions to link macroecology, macroevolution and conservation prioritization. Ecography n/a. https://doi.org/10.1111/ecog.05557

Zizka, A., Silvestro, D., Vitt, P., Knight, T.M., 2021b. Automated conservation assessment of the orchid family with deep learning. Conservation Biology 35, 897–908.

Zizka, A., Andermann, T., & Silvestro, D. (2022). IUCNN – Deep learning approaches to approximate species’ extinction risk. Diversity and Distributions, 28(2), Article 2. https://doi.org/10.1111/ddi.13450

