## Supplementary information for "Modelling the probability of meeting IUCN Red List criteria to support reassessments"

Table S1. Summary of predictors of extinction used in the study.

| Class | Predictor | Description | Source | Rationale |
|---|---|---|---|---|
| Morphology | Body Mass | Log transformed | AVONET (Tobias et al., 2022) | Allometric relationships directly relate body mass to extinction parameters (range size, population density, harvest pressure...) |
| | Beak length nares | Log transformed | AVONET (Tobias et al., 2022) | Beak size relates to diet specialisation which can impact extinction risk |
| | Hand Wing Index | | AVONET (Tobias et al., 2022) | Hand-wing index is a proxy for dispersal abilities, which might affect species' extinction risk |
| Life history | Generation length | Log transformed | BirdLife International, 2022 | Species with long generation length might be more likely to trigger criteria A and C as declines are assessed over a longer timeframe |
| | Clutch size | Log transformed | Tobias and Pigot 2019 | Species with small clutch size might be more likely to decline and thus to be threatened under criteria A and C |
| Behaviour | Migration | 2 classes: Migrant (Full migrant, nomadic, altitudinal migrant), None Migrant (Not a Migrant, Unknown) | BirdLife International, 2022 | Migrant species might be exposed to additional threats in their non-breeding grounds and thus more threatened |
| | Nocturnal activity | 2 classes: Nocturnal or not | | Nocturnal species might be less impacted by human disturbance |
| Ecological | Trophic niche | 4 classes: Herbivore (frugivore, granivore, herbivore terrestrial, herbivore aquatic, nectarivore), Omnivore, Invertivore, Carnivore (scavenger, aquatic predator, vertivore) | AVONET (Tobias et al., 2022) | Some trophic niches are more likely to be affected by some human activities (e.g., insectivorous to pesticide use) |
| | Forest dependency | 4 classes: Non- Forest, Low, Medium, High | BirdLife International, 2022 | Species with high forest dependency might be more sensitive to forest cover loss |
| | Habitat breadth | Number of main habitats listed as suitable in the Red List | BirdLife International, 2022 | Species with high habitat specialisation might be more sensitive to habitat loss |
| Geographical | Insularity | Binary: Insular if > 25% of the range on small islands | Tobias and Pigot 2019 | Isolated species might be more likely to be threatened, especially under criteria B1, B2, and D2 |
| | Range size | Derived from species range<br>Log transformed | BirdLife International, 2022 | Species with small range size might be more likely to be threatened, especially under criteria B1 and B2 |
| Taxonomy | Family | | BirdLife International, 2022 | Extinction might be structured taxonomically |
| Habitat alteration | Canopy density 2018<br>Canopy change 2000-2018 | Raster resolution: 300*300m<br>Operation: median<br>Log transformed | Remelgado and Meyer (Under review) | Loss of forest cover might increase species extinction risk for forest-dependent species |
| | Crop land gains & losses 2003 to 2019<br>Crop land cover 2019 | Resolution: 3,000*3,000m<br>Operation: median<br>Log transformed | Potapov et al. 2022 | Intense agriculture might increase species extinction risk |
| Human disturbance | Per capita Growth Domestic Product | Operation: median between countries of occurrences<br>Log transformed | Worldbank 2021 | Human activities can differ in scope and scale depending on a country's GDP |
| | Population density 2015<br>Trends 2000-2015 | Raster resolution: 1,000*1,000m<br>Operation:<br>- Median<br>- If Pop 2000 > 0, Trends = (Pop 2015 – Pop 2000) / Pop 2000, else if Pop 2000 = 0 and Pop 2015 > 0, Trends = 1, else Trends = 0<br>Log transformed | Global Human Settlement Layer (Schiavina et al., 2019) | Species living in areas of high human density might be more likely to be threatened by human activities |
| | Proportion of rural people 2015 | Raster resolution: 1000*1000m<br>Operation:<br>- If Population 2015 >0, Proportion rural= (Total population 2015 – Total rural 2015) / (Total population 2015), else, Proportion rural = 0 | Global Human Settlement Layer (Schiavina et al., 2019) | Rural and urban human populations might impact species differently (e.g., urban populations via urban development and disturbance, rural populations by agriculture and hunting) |
| | Travel time to cities | Operation: median<br>Log transformed | Weiss et al. 2018 | Species living close to cities might be more likely to be threatened by human activities |
| Climate change | Difference between species Mean annual air temperature and Annual precipitation amount between two periods of time (past and present) | Periods: 1965-1995 and 2005-2014<br>Operation: Median and difference | CHELSA (Karger et al., 2017) from Lucas et al. 2023 | Species exposed to climate change might be more likely to be threatened if sensitive |

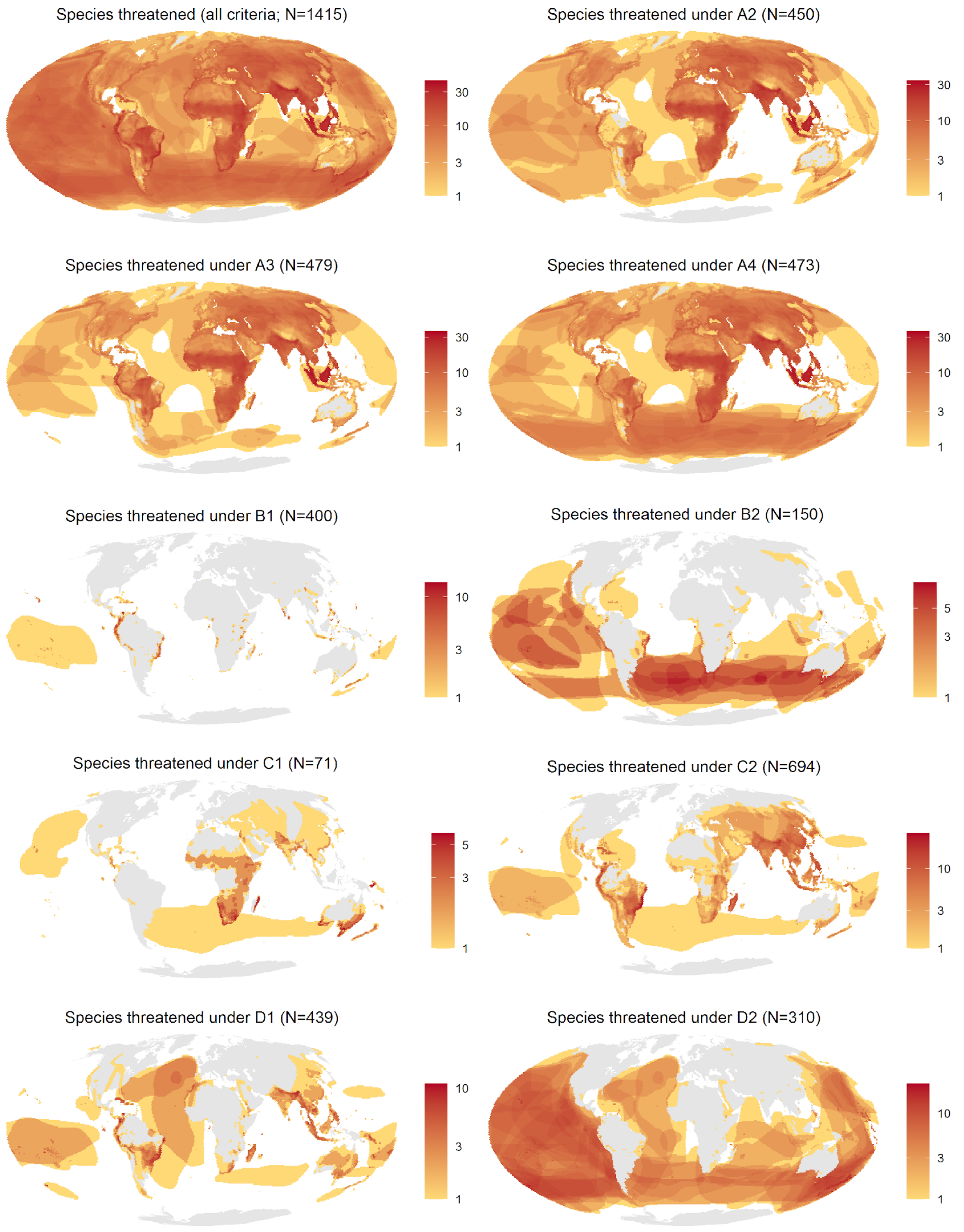


Figure S1: Distribution of threatened species (per BirdLife International 2021) under the nine criteria assessed in the analyses.

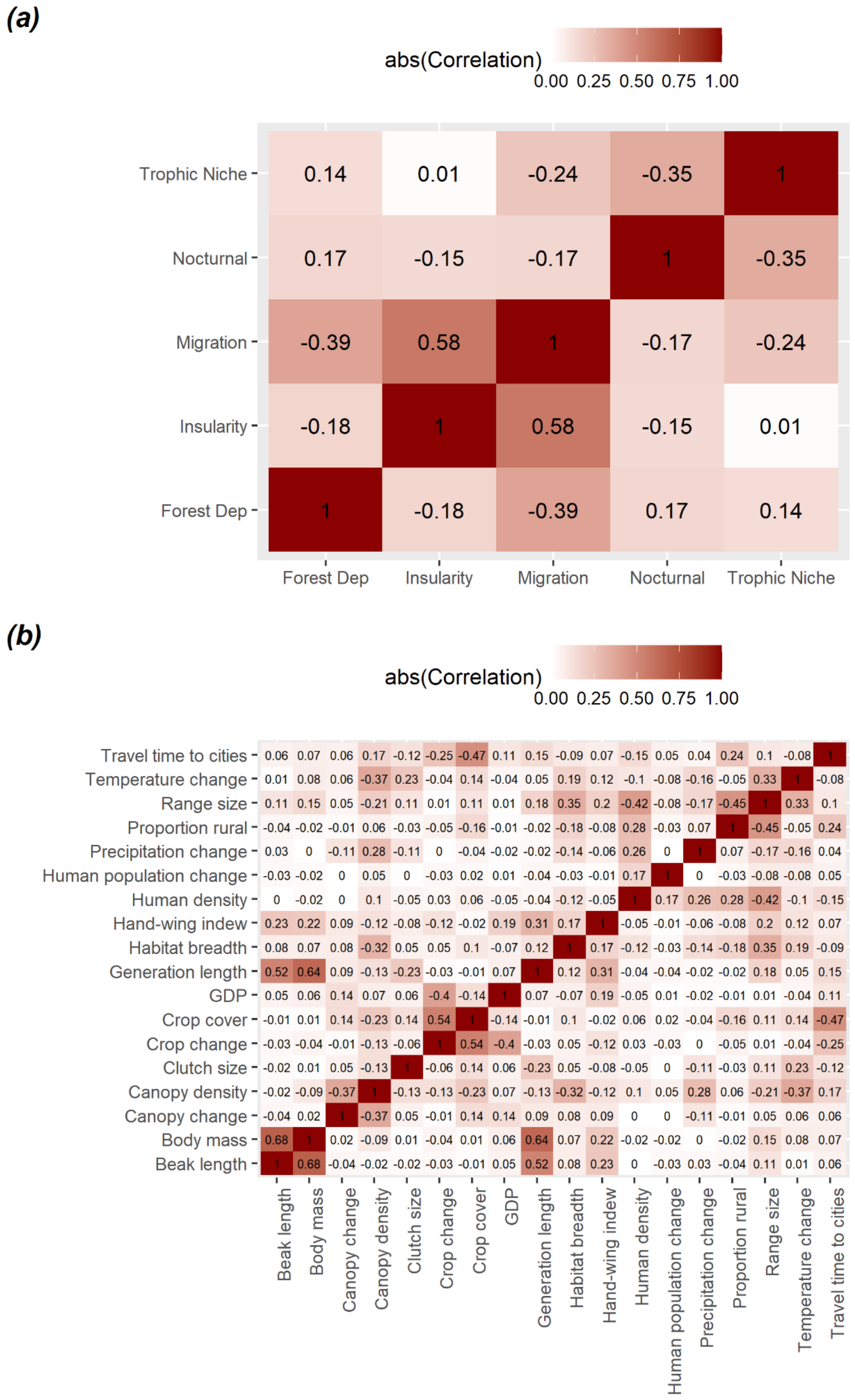


Figure S2: Correlation between (a) qualitative covariates (measured with a Kruskal-gamma test) and (b) quantitative covariates (measured with the Pearson index).

#### Supplementary Methods S1: Choice of climate change data to include in the analyses

In our final models, we used two variables as proxies for climate change: Change in mean annual air temperature (bioclimatic variable 1) and Change in Annual precipitation amount (bioclimatic variable 12), calculated with the methodology presented in the main text. This choice was made to limit the complexity of the model and enable interpretability of the correlation between climate change variables and extinction risk.

We verified that these variables were representative descriptors of species' range climate by running a PCA following Lucas et al. (2023) on the mean values for both time steps (1965-1995 to represent the past climate and 2005-2014 to represent the current climate; (Fig. S3). It shows that the two variables we selected are respectively central in the two main variable clusters from the PCA and are almost orthogonal. We thus considered they were representative of the overall climate differences between species. The spatial distribution of these variables and their change in time is presented in Fig. S4.

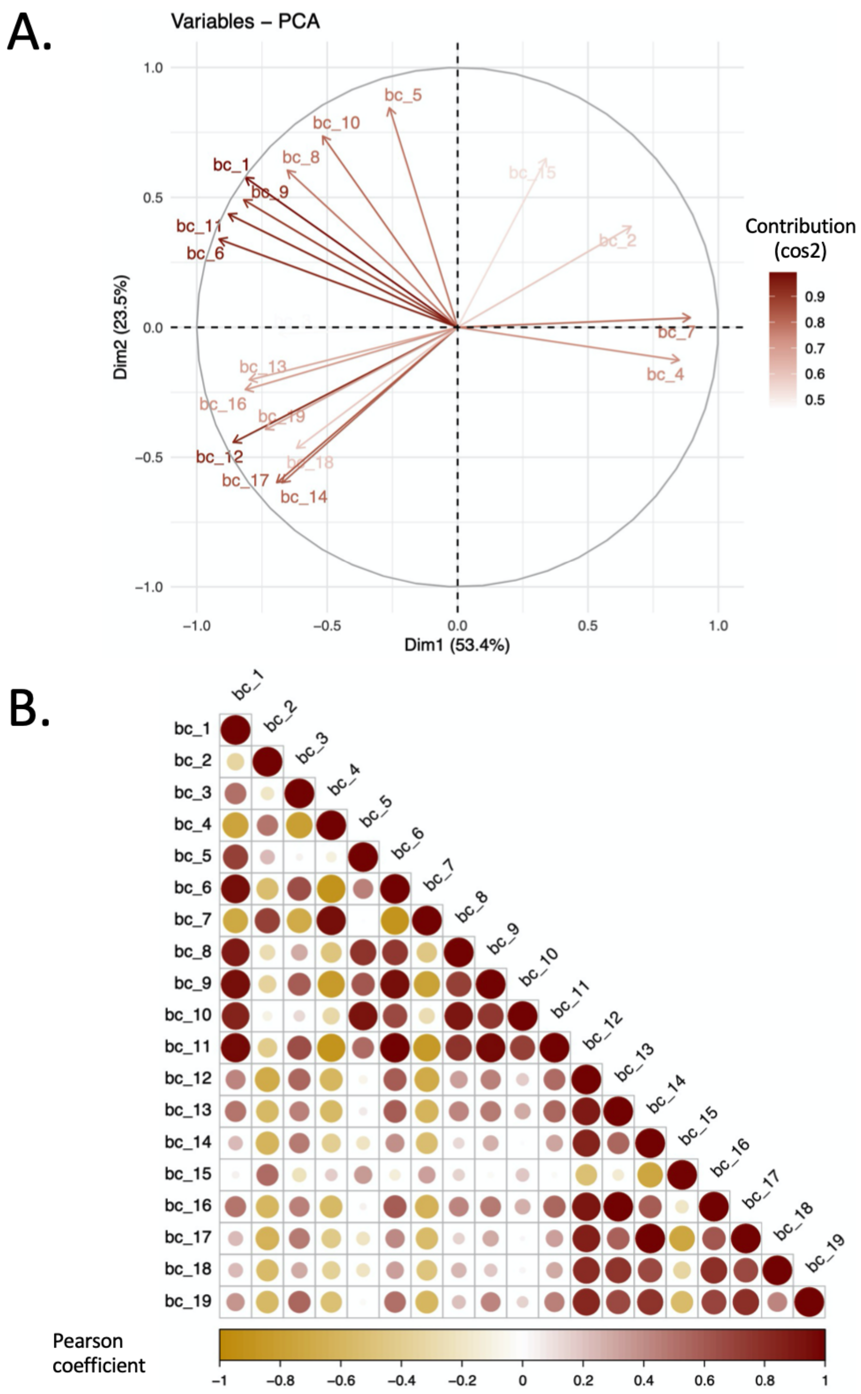


Figure S3. Results of principal component analysis on CHELSA bioclimatic variables. Analyses were performed using the *'factorextra'* package (Kassambara and Mundt, 2020). A: Visualisation of the variables on the first and second axis of the Principal Component Analysis and their contributions (in percentage) of the variables to the principal components. B: Matrix of correlation between bioclimatic variables. Variables: 1. Mean annual air temperature, 2. Mean diurnal air temperature range, 3. Isothermality, 4. Temperature seasonality, 5. Mean daily maximum air temperature of the warmest month, 6. Mean daily minimum air temperature of the coldest month, 7. Annual range of air temperature, 8. Mean daily mean air temperatures of the wettest quarter, 9. Mean daily mean air temperatures of the driest quarter, 10. Mean daily mean air temperatures of the warmest quarter, 11. Mean daily mean air temperatures of the coldest quarter, 12. Annual precipitation amount, 13. Precipitation amount of the wettest month, 14. Precipitation amount of the driest month, 15. Precipitation seasonality, 16. Mean monthly precipitation amount of the wettest quarter, 17. Mean monthly precipitation amount of the driest quarter, 18. Mean monthly precipitation amount of the warmest quarter, 19. Mean monthly precipitation amount of the coldest quarter.

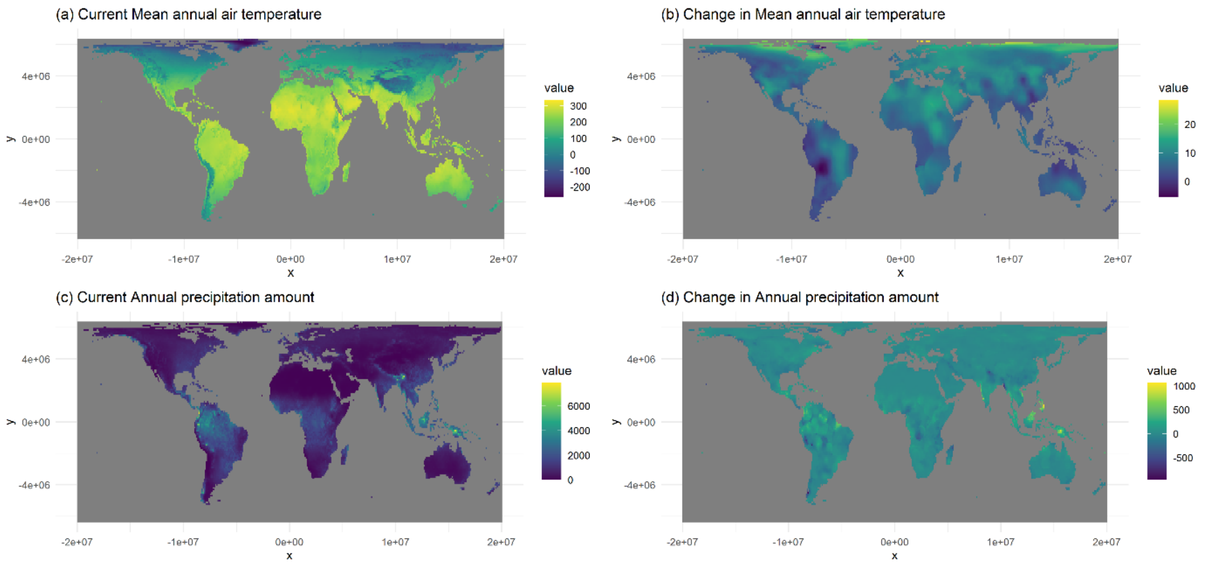

Figure S4: Map of climate and climate change variables kept in the analyses. Temperature variable is 10 times the temperature in °C (i.e., a value of 300 corresponds to 30°C). Precipitation variable is given in kg/m2.

#### Supplementary Methods S2: Test of the proportional odds assumption for the criterion-blind model

Proportional odds logistic regression can be used when there are more than two outcome categories that have an order. It allows us to preserve the ordinal structure of the IUCN Red List categories (LC < NT < VU < EN < CR) considered as the response variable in our models, without necessarily assuming that every category is equidistant. It has been used in several studies to model Red List categories (Lucas et al., 2019; Luiz et al., 2016; Lucas et al., 2023). However, an important underlying assumption is that no input variable should have a disproportionate effect on a specific level of the outcome variable; i.e., the coefficients for each predictor category must be consistent, or have parallel slopes, across all levels of the response. This assumption is known as the proportional odds (PO) assumption.

We thus run several tests to check for the assumption in each model. The first one is the "nominal test" of the package ‘ordinal’ (Christensen, 2019). For a given model, it tests the PO assumption for all variables and when the test is significant, it shows evidence that the PO assumption does not hold for this variable.

To test whether the ‘slope’ of the logistic function of each level of our outcomes was the same, we also then ran stratified binomial models and compared the coefficients of our input variables. We thus created four binomial logistic regression models for the five levels of our outcome variable:

- LC species vs all others (LC)
- LC+NT species vs VU+EN+CR species (NT)
- LC+NT+VU species vs EN+CR species (VU)
- CR species vs all others (EN)

In this way, we modelled the change at each step of our ordinal scale, while keeping the sample size constant.

Fig. S5 shows the results of our test of the PO hypothesis for the criterion-blind model. Although the predictors hand-wing index, forest dependence, and canopy density cover showed strong significance in the nominal test, suggesting that these variables violate the PO assumption, comparison of the coefficients in the binomial models shows no distinct difference (overlapping CIs), suggesting that the estimates of our ordinal scale are not significantly different. However, body mass and range size appear to have significantly different effects between threatened and CR species, respectively. Nevertheless, the mean of the estimates remains below or above 0, indicating that the sign of the relationship is consistent across categories. We obtain similar results for the other models.

In addition, most commonly used modelling alternatives of proportional odds logistic regression (e.g., aggregated binomial classifications or assuming that differences between adjacent risk levels are equivalent) violates assumptions in a much more serious way while leading to a significant loss of information (Senn and Julious, 2009). The latter assumes that the predictors of each threat category have an equal impact on the species' threat level, which is certainly not correct. In contrast, proportional odds logistic regression preserves the structure of the ordinal categorical response variables. The odds ratio, which estimates the degree to which a predictor affects the extinction level, remained consistent across categories (see above; Fig.S5), meaning that even if the OP does not hold, the approximation offered by the OP is still meaningful.These observations led us to keep the proportional odds logistic regression as modelling method for our study.

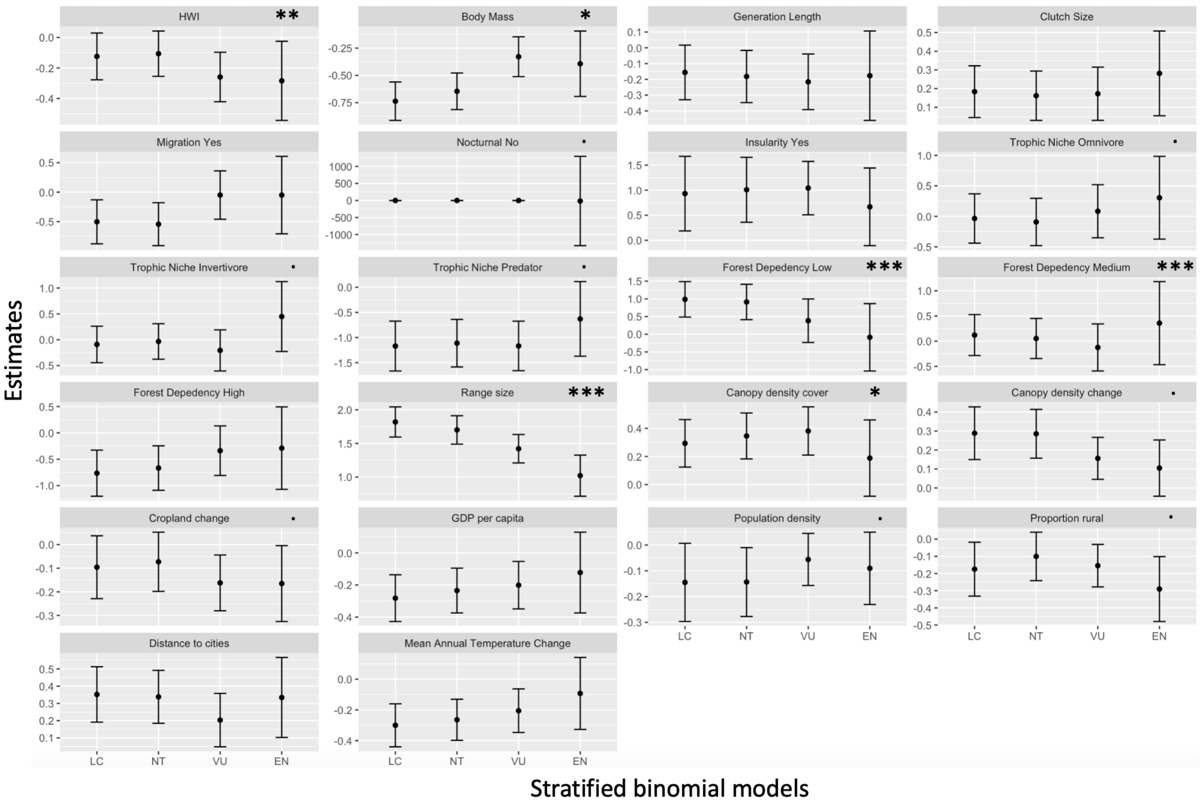


Fig S5: Test of the proportional odds assumption for the criterion-blind model after variable selection. Asterisks represent the significance of the likelihood ratio test resulting from the nominal test and on the x-axis distinguish the stratified binomial models (see text). The y-axis represents the mean of the predictor estimates for each binomial model and the corresponding 95% confidence interval. Asterisks represent the significance of the likelihood ratio test resulting from the nominal test (codes: 0 '***' 0.001 '**' 0.01 '*' 0.05 '.' 0.1 ' ' 1).

### Supplementary Results

Table S2. Models' performances under block-taxonomic validation, as displayed in Fig. 2. For each model, the table provides the number of species used to train the model ('n'), the proportion of threatened species according to that specific criteria ('% Threatened species'), the model sensitivity, specificity and TSS.

| Models | n | % Threatened species | Sensitivity | Specificity | TSS |
|---|---|---|---|---|---|
| Criterion-blind | 8999 | 11.78 | 0.76 | 0.82 | 0.58 |
| A2 | 7844 | 4.83 | 0.76 | 0.81 | 0.57 |
| A3 | 7923 | 5.10 | 0.70 | 0.79 | 0.50 |
| A4 | 7873 | 4.95 | 0.74 | 0.81 | 0.55 |
| B1 | 7575 | 3.83 | 0.96 | 0.91 | 0.87 |
| B2 | | 1.39 | 0.80 | 0.93 | 0.73 |

| | | | | | |
|---|---|---|---|---|---|
| C1 | 7336 | 0.78 | 0.86 | 0.85 | 0.71 |
| C2 | 7902 | 6.82 | 0.82 | 0.86 | 0.69 |
| D1 | 7452 | 3.84 | 0.87 | 0.90 | 0.78 |
| D2 | 7343 | 2.46 | 0.88 | 0.92 | 0.79 |
| Criterion-specific | 8999 | 11.78 | 0.83 | 0.69 | 0.50 |

Table S3. Models' performances under block-taxonomic validation when considering missing data on the category triggered by a species under a specific criterion (see Fig. 1) as LC. Column names as in Table S2.

| Models | n | % Threatened species | Sensitivity | Specificity | TSS |
|---|---|---|---|---|---|
| Criterion-blind | 8999 | 11.8 | 0.76 | 0.82 | 0.58 |
| A2 | 8999 | 4.21 | 0.69 | 0.79 | 0.48 |
| A3 | 8999 | 4.49 | 0.62 | 0.78 | 0.40 |
| A4 | 8999 | 4.33 | 0.68 | 0.79 | 0.47 |
| B1 | 8999 | 3.22 | 0.95 | 0.89 | 0.84 |
| B2 | 8999 | 1.12 | 0.71 | 0.90 | 0.62 |
| C1 | 8999 | 0.63 | 0.81 | 0.81 | 0.62 |
| C2 | 8999 | 5.99 | 0.78 | 0.84 | 0.62 |
| D1 | 8999 | 3.18 | 0.85 | 0.88 | 0.72 |
| D2 | 8999 | 2.01 | 0.83 | 0.88 | 0.71 |
| Criterion-specific | 8999 | 11.8 | 0.87 | 0.68 | 0.55 |

Table S4. Models' performances under block-taxonomic validation at the category level.

| Model | Sensitivity | Specificity | TSS | Accuracy | | | | |
|---|---|---|---|---|---|---|---|---|
| | | | | LC | NT | VU | EN | CR |
| Criterion-blind | 0.40 | 0.77 | 0.18 | 0.86 | 0.00 | 0.51 | 0.27 | 0.22 |
| A2 | 0.46 | 0.79 | 0.25 | 0.82 | 0.00 | 0.63 | 0.08 | 0.00 |
| A3 | 0.47 | 0.77 | 0.24 | 0.81 | 0.00 | 0.61 | 0.12 | 0.00 |
| A4 | 0.46 | 0.79 | 0.25 | 0.83 | 0.00 | 0.62 | 0.16 | 0.04 |
| B1 | 0.66 | 0.91 | 0.57 | 0.93 | 0.00 | 0.71 | 0.65 | 0.31 |
| B2 | 0.47 | 0.93 | 0.39 | 0.92 | 0.00 | 0.19 | 0.67 | 0.00 |
| C1 | 0.65 | 0.84 | 0.49 | 0.86 | 0.00 | 0.76 | 0.00 | 0.40 |
| C2 | 0.47 | 0.85 | 0.32 | 0.88 | 0.00 | 0.58 | 0.28 | 0.38 |
| D1 | 0.49 | 0.91 | 0.40 | 0.91 | 0.00 | 0.61 | 0.33 | 0.27 |
| D2 | 0.88 | 0.92 | 0.79 | 0.92 | 0.00 | 0.88 | 0.00 | 0.00 |
| Criterion-specific | 0.49 | 0.63 | 0.13 | 0.70 | 0.00 | 0.59 | 0.38 | 0.33 |
