## Supplementary material for "Modelling the probability of meeting IUCN Red List criteria to support reassessments": Criterion specific predictions: READ_ME.rtf

We predicted Red List category for each criterion using ordinal regression models (see Methods). 
LC species were considered LC under every criterion, as recommended by the Red List assessment process. 

The table includes the following columns: 
	- Family: species family after taxonomic corrections (see Methods). 
	- Species: species scientific name after taxonomic corrections (see Methods). 
	- RedList_category: Red List category in the last published assessment.
	- Criterion_blind_predicted: Predicted Red List category when actual Red List category is used as  response variable and contrasted to a set of extinction predictors (as is the case in classical extinction risk analysis). 
	- A1: category met under criterion A1 in the last published assessment.
 	- A2: category met under criterion A2 in the last published assessment.
	- A2_predicted : category that our criterion-specific model predicted for the criterion A2.
	- A3: category met under criterion A3 in the last published assessment.
	- A3_predicted : category that our criterion-specific model predicted for the criterion A3.	- A4: category met under criterion A4 in the last published assessment.
	- A4_predicted : category that our criterion-specific model predicted for the criterion A4.
	- B1: category met under criterion B1 in the last published assessment.
	- B1_predicted : category that our criterion-specific model predicted for the criterion B1.
	- B2: category met under criterion B2 in the last published assessment.
	- B2_predicted : category that our criterion-specific model predicted for the criterion B2
	- C1: category met under criterion C1 in the last published assessment.
	- C1_predicted : category that our criterion-specific model predicted for the criterion C1.
	- C2: category met under criterion C2 in the last published assessment.
	- C2_predicted : category that our criterion-specific model predicted for the criterion C2.
	- D1: category met under criterion D1 in the last published assessment.
	- D1_predicted : category that our criterion-specific model predicted for the criterion D1.	- D2: category met under criterion D2 in the last published assessment.
	- D2_predicted : category that our criterion-specific model predicted for the criterion D2.
	- Criterion_spe_combined: final category based on the highest predicted extinction risk among each criterion-specific model (i.e., threatened if at least one of the criteria was predicted to be triggered) as done during Red List assessments. 
